# Biallelic Mutations in *LRRC56* encoding a protein associated with intraflagellar transport, cause mucociliary clearance and laterality defects

**DOI:** 10.1101/288852

**Authors:** Serge Bonnefoy, Christopher M. Watson, Kristin D. Kernohan, Moara Lemos, Sebastian Hutchinson, James A. Poulter, Laura A. Crinnion, Chris O’Callaghan, Robert A. Hirst, Andrew Rutman, Lijia Huang, Taila Hartley, David Grynspan, Eduardo Moya, Chunmei Li, Ian M. Carr, David T. Bonthron, Michel Leroux, Care4Rare Canada Consortium, Kym M. Boycott, Philippe Bastin, Eamonn G. Sheridan

## Abstract

Defective motile cilia are responsible for a group of heterogeneous genetic conditions characterised by dysfunction of the apparatus responsible for generating fluid flows. Primary ciliary dyskinesia (PCD) is the prototype for such disorders and presents with impaired pulmonary mucus clearance, susceptibility to chronic recurrent respiratory infections, male infertility and laterality defects in about 50 % of patients. Here we report biallelic variants in *LRRC56* (also known as ODA8), identified in two unrelated consanguineous families. The phenotype comprises laterality defects and chronic pulmonary infections. High speed video microscopy of cultured patient epithelial cells showed severely dyskinetic cilia, but no obvious ultra-structural abnormalities on routine transmission electron microscopy (TEM). Further investigation revealed that LRRC56 interacts with the intraflagellar transport (IFT) protein IFT88. The link to IFT was interrogated in *Trypanosoma brucei.* In this protist, LRRC56 is recruited to the cilium during axoneme construction, where it co-localises with IFT trains and facilitates the addition of dynein arms to the distal end of the flagellum. In *T. brucei* carrying LRRC56 null mutations, or a mutation (p.Leu259Pro) corresponding to the p.Leu140Pro variant seen in one of the affected families, we observed abnormal ciliary beat patterns and an absence of outer dynein arms restricted to the distal portion of the axoneme. Together, our findings confirm that deleterious variants in *LRRC56* result in a human disease, and suggest this protein has a likely role in dynein transport during cilia assembly that is evolutionarily important for cilia motility.

## INTRODUCTION

Cilia are highly conserved eukaryotic organelles that are classified into motile and non-motile forms. Defective motile cilia underlie the pathophysiology of patients with impaired mucociliary clearance, which increases susceptibility to respiratory complications, including sinusitis, bronchitis, pneumonia, and otitis media^1^. Chronic infections frequently lead to progressive pulmonary damage and bronchiectasis. Spermatozoal dysmotility in affected men causes infertility^1^. Approximately half of all patients with the prototypical disease of motile cilia, Primary Ciliary Dyskinesia ([MIM: PS244400]), display laterality defects varying from partial to complete *situs inversus*, a consequence of dysfunctional embryonic nodal cilia^1^. At least 36 genes currently account for the heterogeneous genetic disorder PCD, which displays mainly autosomal recessive inheritance and is characterised by cilia dyskinesis and structural defects observed by TEM of a nasal biopsy. However this is not always the case. Variants in *CCNO* ([MIM: 607752]) cause a reduction in the generation of motile cilia and result in a similar phenotype ([CILD29 MIM:615872]), except that it is not associated with laterality defects^2^. Mutations in *DNAH11* ([MIM: 603339]) result in CILD7 ([MIM: 611884]) due to an abnormally rapid ciliary beat frequency without a discernible structural defect^3^. These data highlight the molecular complexity underlying the formation and function of cilia^1^.

Motile cilia and flagella, a hallmark of eukaryotes, display remarkable structural and molecular conservation^4^. Most motile cilia exhibit a 9+2 configuration—a pair of single microtubules surrounded by 9 peripheral doublets. Connected to each doublet of peripheral microtubules is an inner and outer dynein arm, consisting of multiple dynein chains that provide ATPase-mediated energy for movement^1^. Dynein arms are preassembled in the cytosol and are transported to an assembly site at the distal end of the growing axoneme. This requires intraflagellar transport (IFT)^5,6 7^, an evolutionary conserved bidirectional transport system that delivers axoneme building blocks^8^,^9^ such as tubulin, to the flagellar tip^10^.

Here, we report biallelic variants in *LRRC56* in patients with bronchiectasis and laterality defects, and show human LRRC56 interacts with the IFT subunit, IFT88. Functional studies using *Trypanosoma brucei* reveal that LRCC56 is recruited during axoneme construction, associates with IFT trains, and is required for the addition of dynein arms to the distal segment of the flagellum. Our findings add *LRRC56* to an expanding list of genes whose disruption results in an atypical ciliary phenotype and reveal a mechanism whereby disruption of LRRC56 leads to defective IFT-dependent targeting of dynein to cilia, and loss of ciliary motility.

## MATERIAL AND METHODS

### Subject evaluation

Two unrelated families were independently ascertained with features suggestive of a ciliopathy (Figure 1A). Family 1 consisted of a single affected female with PCD, whose parents are first cousins of UK Pakistani ethnicity. She presented with chronic chest infections; nasal biopsy was obtained and respiratory epithelial cultures prepared. These were investigated by transmission electron microscopy (TEM) and high-speed video microscopy. Family 2 consisted of two affected individuals, the offspring of first cousin consanguineous parents from Kuwait, ascertained during pregnancy to have lethal congenital heart disease. Both pregnancies were terminated and post mortem pathological investigations were performed. The families provided signed informed consent to participate in studies approved by the Leeds East Research Ethics Committee (07/H1306/113; Family 1) and the Review Ethics Board of the Children’s Hospital of Eastern Ontario (11/04E; Family2).

**Figure 1.**
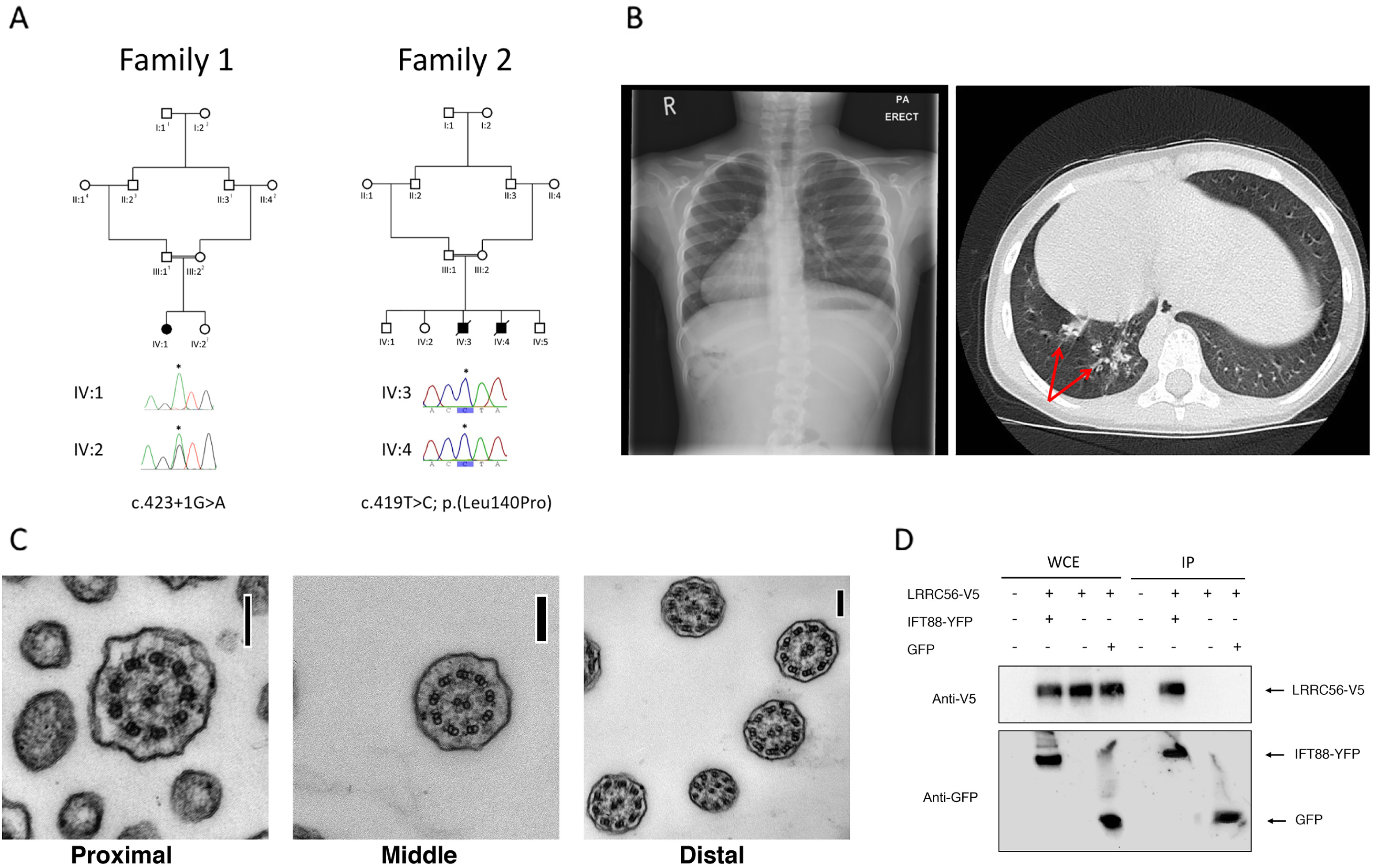
LRRC56 mutations cause PCD and laterality defects. (A) The homozygous splice-site mutation (c.423+1G>A, NM_198075.3) disrupts the evolutionarily conserved canonical donor site in Family 1 individual IV:1. The unaffected sibling IV:2 is heterozygous for the mutation. The homozygous missense mutation (c.419T>C, NM_198075.3) was identified in the two affected siblings V:3 and V:4 from Family 2. Consistent with autosomal-recessive inheritance, the mutations described were detected in a heterozygous state in the unaffected parents (data not shown). (B) The Family 1 proband (IV:1) had dextrocardia, documented by chest X-ray (left panel). High-resolution axial computed tomography of the thorax in the same patient demonstrates bronchiectasis, typical of PCD. Dextrocardia is also visible (CT-scan; right panel). (C) Cross section through the axoneme from cultured respiratory cells from Family 1 individual IV:1. The position of the section is indicated, bar = 100 nm. Normal axonemal structure is visible, with intact dynein arms. (D) Recombinant human LRRC56 and intraflagellar transport protein IFT88 interact in vitro. HEK293 cells were co-transfected with plasmids encoding human LRRC56 and IFT88 tagged with V5 and YFP respectively (1.5µg of each). After 48 hours, immunoprecipitation (IP) was performed with transfected and untransfected cells using Cell-TRAP magnetic beads bound to an anti-GFP antibody fragment. Protein from input whole cell extracts (WCE, left) and immunoprecipitated proteins (IP, right) were blotted using anti-V5 or anti-GFP. A β-actin control is also shown.

### Genetic Analysis

In view of the consanguinity in both families, genetic analysis was performed under an autosomal recessive model. Whole exome sequencing of genomic DNA was performed at the University of Leeds and the Children’s Hospital of Eastern Ontario. Target enrichment was performed, following manufacturer’s protocols, using SureSelect hybridization capture reagents with v5 and v4 Human All Exon probes for Family 1 and 2, respectively (Agilent Technologies). Enriched library preparations were sequenced on either HiSeq 2000/2500 platforms (Illumina) using paired end 100-bp reads. Bioinformatic data processing was performed as previously described, with all variants being reported against human reference genome build hg19^11,12^. Genomic regions corresponding to runs of autozygosity were identified from pipeline-produced variant call format (VCF) files using the tool AgileMultiIdeogram (see Web Resources). These were subsequently used to filter Alamut Batch-annotated variant reports. Additional filtering criteria included the exclusion of common variants (those with a minor allele frequency ≥1%) represented in the NHLBI Exome Variant Server (EVS) or an in-house cohort of >1,500 control exomes and the exclusion of genes with biallelic functional variants reported to the Exome Aggregation Consortium (ExAC). Pathogenic variants and segregation in the families were confirmed using Sanger sequencing with an ABI3130xl. Primer sequences and thermocycling conditions are available upon request.

### RNA splicing assay

Total RNA was extracted from peripheral blood of the affected proband in Family 1 using the QIAamp RNA blood mini kit (Qiagen). A gene-specific primer spanning the boundary of *LRRC56* exons 10 and 11 (NM_198075.3) (dCTTGGCCAGCACCATGGGTGAG) was used to perform first-strand cDNA synthesis with a SuperScript^TM^ II RT kit (Life Technologies). Two PCR amplicons were generated from the resulting cDNA using an exon 6 forward primer (dCAACCTGGACCAACTGAAGC) combined with either an exon 8 (dCCTCCAGGGTGAGCATGG) or exon 10 (dCCAGGTCCTCAGAAAGCAGG) reverse primer. PCR products were used to create Illumina-compatible sequencing libraries with NEBNext^®^ Ultra^TM^ reagents (New England Biolabs) and sequenced on an Illumina MiSeq using a paired-end 150bp read length configuration.

### Co-IP experiments

Human *LRRC56* was amplified from an untagged image clone (SC123392, Origene Technologies), inserted into the pDONR201 Gateway cloning vector (Invitrogen) and subsequently pDEST40 destination vector according to the manufacturer’s instructions. The human IFT88-eYFP construct was provided by Professor Colin Johnson^13^. Both plasmids were sequenced and maxi-prepped prior to transfection. HEK293 cells were co-transfected with 1.5µg of each plasmid using Lipofectamine 2000 (Invitrogen). Cells were lysed 48 h post transfection, using NP40 cell lysis buffer, containing 1x protease and phosphatase inhibitors, and protein extracted as per standard protocols. Immunoprecipitation (IP) of eYFP was performed with 1000 µg of both transfected and untransfected cell extracts using the GFP-Trap^®^_M kit (Chromotek) according to the manufacturer’s instructions. Whole cell extracts (WCE) and immunoprecipitates (IP) were blotted using mouse anti-V5 (Invitrogen, 1/2,000) or mouse anti-GFP (Invitrogen, 1/1,000). A secondary goat anti-mouse HRP (Dako) was used at 1/10,000 and detected using the SuperSignal West Femto kit (Thermo Fisher Scientific).

### Ciliary function measurements

In IV:1, Family 1, ciliary beat frequency (CBF) was measured using a digital high-speed video imaging system, as described previously^14,15^. Ciliary beat pattern was investigated in three different planes: a sideways profile, beating directly toward the observer and from directly above. Dyskinesia was defined as an abnormal beat pattern that included reduced beat amplitude, stiff beat pattern, flickering motion or a twitching motion. Ciliary beat pattern is associated with specific ultrastructural defects in primary ciliary dyskinesia. Dyskinesia index was calculated as the percentage of dyskinetic cilia within the sample (number of dyskinetic readings/total number of readings for sample ×100). All measurements were taken with the solution temperature between 36.5 and 37.5° C and the pH between 7.35 and 7.45. Normal ranges for CBF and percentage dyskinetic cells are 8-17Hz and 4-49% respectively as previously reported^16^.

### Air liquid interface cell culture

The modified method we used has previously been described^17^. Briefly, nasal brush biopsy samples were grown on collagen (0.1%, Vitrogen, Netherlands) coated tissue culture trays (12 well) in Bronchial Epithelial Growth Media (BEGM, Lonza, USA) for 2-7 days. The confluent unciliated basal cells were expanded into collagen-coated 80 cm^2^ flasks and the BEGM was replaced every 2-3 days. The basal cells were then seeded at approximately 1-3 × 10^5^ cells per cm^2^ on a collagen coated 12mm diameter transwell clear insert (Costar, Corning, USA) under BEGM for 2 days. After confluency, the basal cell monolayer was fed on the basolateral side only with ALI-media (50% BEGM and 50% Hi-glucose minimal essential medium containing 100 nM retinoic acid). The media was exchanged every 2-3 days and the apical surface liquid was removed by gentle washing with phosphate buffered saline when required. When cilia were observed on the cultures they were physically removed from the transwell insert by gentle scraping with a spatula and washing with 1ml of HEPES (20 mM) buffered medium 199 containing penicillin (50 µg/ml), streptomycin (50 µg/ml) and Fungizone (1 µg/ml). The recovered ciliated epithelium was then dissociated by gentle pipetting and 100µl of the cell suspension was examined inside a chamber slide and ciliary beat frequency and pattern assessed as described above ^14,15^. The remaining 900 µl was fixed in 4% glutaraldehyde for transmission electron microscopy (TEM) analysis of cilia and ciliated epithelium.

### TEM of Human Samples

TEM was performed as previously described^18^, using a Jeol 1200 instrument. For TEM, the ciliated cultures were fixed with glutaraldehyde (4%w/v) in Sorensen’s phosphate buffered (pH 7.4). After post-fixation in osmium tetroxide (1%w/v), samples were dehydrated through graded ethanol series and immersed in hexamethyldisilazane. Sections were cut at 90nm, with cross-sections categorised for height in the cilium using histological parameters. The bottom (2µm) of the axoneme cross-sections were located because they were associated with microvilli. The middle (2µm) cross sections were identified by wide cross-sections with the outer dynein (ODA) away from the axoneme membrane (with no microvilli present). The top 1um of the cilia was represented by cross-sections which have a slightly smaller diameter compared to the middle sections. In addition, the axonemal membrane was tightly wrapped to the microtubules and the ODA. The tips (0.6µm) appear as thin cross sections and have no dynein arms.

### Trypanosome cell culture

All cloned cell lines used for this work were derivatives of *T. brucei* strain Lister 427 and were cultured in SDM79 medium supplemented with hemin and 10 % fetal calf serum^19^. Cell lines *IFT88^RNAi^* (targeting an essential protein for anterograde IFT)^20^, *IFT140^RNAi^* (essential protein for retrograde IFT)^21^, *DNAI1^RNAi^* (component of the dynein arm)^22^, and *ODA7^RNAi^* (cytoplasmic assembly machinery of the dynein arm)^23^ have previously been described. They all express double-stranded RNA under the control of tetracycline-inducible T7 promoters^24,25^.

### Expression of endogenous LRRC56 fused to YFP

Endogenous tagging of the *T. brucei LRRC56* gene was carried out by direct integration in the *LRRC56* gene using the p2675LRRC56 plasmid. This vector is derived from the p2675 plasmid and contains a 1495bp fragment of the trypanosome *LRRC56* gene sequence (1-1495) downstream of the *YFP* gene^26^. Transfection was achieved by nucleofection of *T. brucei* cells using program X-0014 of the AMAXA Nucleofector*®* apparatus (Lonza) as previously described ^27^, with 10 µg plasmid linearized with AfeI in the target *LRRC56* gene sequence, for homologous recombination with the target allele. The mutant allele was obtained following T776/C base substitution (Genecust Europe, Luxembourg) to mutate leucine 259 to proline in the p2675TbLRRC56. The resulting p2675LRRC56L259P plasmid was linearized with NheI prior to nucleofection. As a result, the expression of the YFP fluorescent fusion protein is under the control of the endogenous 3′ untranslated region of the *LRRC56* gene resulting in a similar expression level as the wild-type allele since most gene expression regulation is determined by 3′UTR sequences^28^. Transgenic cell lines were obtained following puromycin selection and cloning by serial dilution.

### *LRRC56* gene deletion or replacement

Sequential *LRRC56* gene replacement in *T. brucei* was used to obtain *lrrc56-/-* cells. Sequences containing either the puromycin (*PURO*) or the blasticidin (*BLA*) drug resistance gene flanked by 300 bp long upstream and downstream regions of the *LRRC56* gene were synthesized and cloned in a pUC57 plasmid (Genecust Europe, Luxembourg). Amplicons were generated by PCR with primers P1: dTTTGAAGGTGCTGTGTGAGGG and P2 dAGGTAGAGGGAGGCGTTGAG (Eurogentec) which anneal to the sequences 300 bp upstream of the *LRRC56* start codon and downstream of *LRRC56* stop codon respectively. Prior to nucleofection PCR fragments containing either the blasticidin (*BLA*) resistance gene or the G418 neomycin (*NEO*) resistance gene were purified using Nucleospin gel and PCR clean-up kit (Macherey Nagel). Single allele knockout cells were obtained following nucleofection with the *BLA* amplicon containing *LRRC56* flanking sequences, blasticidin selection and cloning. *LRRC56* allele deletion was confirmed using PCR with 5′UTR *LRRC56*-specific primer P3 dTCACCATCACGCCCTTTTGT and *BLA*-specific primer P4 dCTGGCGACGCTGTAGTCTT. Replacement of the remaining *LRRC56* allele was performed with either the *YFP::LRRC56* gene or the mutated *YFP::LRRC56L259P* gene in this single knockout cell line using linearized p2675TbLRRC56 or p2675TbLRRC56L259P plasmid and puromycin selection as described above. Double *LRRC56* knockout was achieved following nucleofection of the cell line with a single allele knockout carrying the mutated *YFP::LRRC56L259P* gene with *NEO* amplicon containing *LRRC56* flanking sequences and validation obtained following PCR analysis of G418-resistant cells with LRRC56-specific primers P5 dCCGTAGCATCATCCGAGACC and P6 dACTATTTGCGTCAGGTGGCA. Primers amplifying a 1511-bp region of the unrelated aquaporin 1 gene served as positive controls. For whole-genome sequencing, genomic DNA was extracted using the Qiagen DNeasy kit. Short insert Illumina sequencing libraries were constructed and sequenced at the Beijing Genomics Institute, generating approximately 6.6 million 100-bp paired end reads. Reads were aligned to the *T. brucei* TREU927 genome (TriTrypDB release 35)^29^ using bowtie2 in very-sensitive-local alignment mode, with an 88.6% alignment rate^29,30^. Alignment files were sorted, merged and indexed using Samtools^31^. Aligned reads were visualised and analysed, including counting reads per CDS using the Artemis pathogen sequence browser^32^.

### Indirect immunofluorescence assay (IFA)

Cultured trypanosomes were washed twice in SDM79 medium without serum and spread directly on poly-L-lysine coated slides (Thermoscientific, Menzel-Gläser) before fixation. For methanol fixation, cells were air-dried and fixed in methanol at −20°C for 5 min followed by a rehydration step in PBS for 15 min. For paraformaldehyde-methanol fixation, cells were left for 10 min to settle on poly-L-lysine coated slides. Adhered cells were washed briefly in PBS before being incubated for 5 min at room temperature with a 4% PFA solution in PBS at pH 7 and fixed with methanol at −20°C for an additional 5 min followed by a rehydration step in PBS for 15 min. For immunodetection, slides were incubated for 1 h at 37°C with the appropriate dilution of the first antibody in 0.1% BSA in PBS; mAb25 recognises the axonemal protein TbSAXO1^33^; a monoclonal antibody against the IFT-B protein IFT172^21^, and a mouse polyclonal antiserum against DNAI1 ^23^. YFP::LRRC56 was observed upon fixation by immunofluorescence using a 1:500 dilution of a rabbit anti-GFP antibody that detects YFP (Life Technologies). After several 5 min washes in PBS, species and subclass-specific secondary antibodies coupled to the appropriate fluorochrome (Alexa 488, Cy3 or Cy5, Jackson ImmunoResearch) were diluted 1/400 in PBS containing 0.1% BSA and were applied for 1 h at 37°C. After washing in PBS as indicated above, cells were stained with a 1 µg/ml solution of the DNA-dye DAPI (Roche) and mounted with Slowfade antifade reagent (Invitrogen). Slides were either stored at −20°C or immediately observed with a DMI4000 microscope (Leica) with a 100X objective (NA 1.4) using a Hamamatsu ORCA-03G camera with an EL6000 (Leica) as light source. Image acquisition was performed using Micro-manager software and images were analysed using ImageJ (National Institutes of Health, Bethesda, MD)

### Motility analyses

Volumes of 250µl at 5×10^6^ trypanosomes/ml in warm medium were distributed in 24 well plates. Samples were analysed under 10X objective of a DM IL LED microscope (Leica) coupled to a DFC3000G camera (Leica). Movies were recorded (200 frames, 50 ms of exposure) using LASX Leica software, converted to .avi files and analysed with the medeaLAB CASA Tracking v.5.5 software (medea AV GmbH) ^34^.

### TEM analysis of trypanosome samples

Cells were fixed with 2.5% glutaraldehyde directly in the suspension culture for 10 min and then treated with 4% paraformaldehyde, 2.5% glutaraldehyde in 0.1M cacodylate buffer (pH 7.2) for 1h. Cells were post-fixed for 30 min (in the dark) in 1 % osmium tetroxide (OsO4), in 0.1 M cacodylate buffer (pH 7.2), washed and incubated in 2% uranyl acetate for 1h at room temperature. Samples were washed, dehydrated in a series of acetone solutions of ascending concentrations, and embedded in Polybed 812 resin. Cytoskeletons were extracted by treating cells with Nonidet 1% with protease inhibitor cocktail (Sigma, P8340) diluted in PBS for 20 min, washed in PHEM 0.1M pH7.2 for 10min. Fixation was performed in 2.5% glutaraldehyde, 4% paraformaldehyde, and 0.5% tannic acid in 0.1M cacodylate buffer (pH 7.2). Cells were post-fixed, incubated in uranyl acetate, dehydrated and infiltrated as cited above. Ultrathin sections were stained with uranyl acetate and lead citrate.

## RESULTS

### Clinical characterization of families

Individual IV:1, Family 1 was born at term by normal vaginal there was no family history of note. After 36 hours, she developed tachypnoea, which was managed with head box oxygen. She was discharged home on day 5 of life. She developed rhinorrheoa and experienced several episodes of documented respiratory tract infection. She suffered from a chronic cough and recurrent middle ear infections. At age 18 months, she developed pneumonia. Chest X-ray and CT scan revealed bronchiectasis and dextrocardia (Figure 1B), suggesting a clinical diagnosis of PCD. However, TEM analysis of nasal biopsy samples along different segments of the cilia (tip, middle and base) showed apparently intact dynein arms (Figure 1C). Direct high-speed videomicroscopy analysis of biopsy material revealed a ciliary beat frequency of 10.87 Hz (10.4 Hz-11.34 Hz) and particulate clearance was observed (Supplementary Video 1 and 2). Although cell culture at an air-liquid interface produced a healthy ciliated epithelium, the ciliary beat pattern was in fact dyskinetic (19% twitching, 33% stiff, 48% static), with a beat frequency of 5.39 Hz (4.29 Hz-6.49 Hz) (Supplementary Video 3 and 4).

Family 2 were from Kuwait and included two affected foetuses. Both pregnancies were terminated because the fetuses were found to have lethal congenital heart disease. Autopsy revealed similar findings in both patients, which included *situs inversus* of the thoracic and abdominal organs, with a complex congenital heart malformation characterized by double outlet right ventricle (data not shown).

### Genetic analyses revealed mutations in *LRRC56*

Autozygosity mapping identified 46 and 33 regions of homozygosity in each proband from Family 1 and 2 respectively (Table S1). Variant filtering, using the autozygous intervals, and public/in-house databases (to eliminate variants with minor allele frequency > 1%) identified a single homozygous variant in each family, located in the same gene, *LRRC56* (NM_198075.3: Family 1 c.423+1G>A; Family 2 c.419T>C, p.(Leu140Pro)). Neither variant is recorded in ExAC. A search of an ethnically matched control exome cohort revealed a single heterozygous carrier of the c.423+1G>A mutation among 1,541 normal subjects. Sanger sequencing confirmed the variants as well as segregation in both families. c.423+1G>A is predicted to abrogate the intron 7 donor site; high-throughput sequencing analysis of RT-PCR products confirmed that the *LRRC56* variant is aberrantly spliced and not predicted to encode a functional protein (Figure S1). The missense variant c.419T>C, p.(Leu140Pro) affects a highly conserved residue (Figure S2), predicted to be deleterious by SIFT (0)^35^, Polyphen (1.0)^36^, and CADD (15.72) scores^37^.

### LRRC56 protein is conserved in eukaryotes with motile cilia constructed by intraflagellar transport (IFT)

The human *LRRC56* gene encodes a 542 amino acid protein determined from transcriptomic studies and immunohistochemistry to be expressed in many organs, including lung and respiratory epithelial cells, but mostly in testis and pituitary gland^38^. The protein contains 5 leucine-rich repeat domains conserved across species whose motile cilia are assembled by intraflagellar transport (IFT)^39^ *LRRC56* is the human orthologue of the *Chlamydomonas reinhardtii* gene *ODA8*, which is thought to play a role in the maturation and/or transport of outer dynein arm (ODA)complexes during flagellum assembly, and has a biochemical distribution similar to IFT proteins^39^. ODA8 is proposed to function together with two other proteins termed ODA5 and ODA10 to form an accessory complex involved in ODA docking to microtubules^39^. However, evidence for a physical interaction is lacking, and the exact role of ODA8 remains to be defined. To evaluate a possible association of LRRC56 with the multi-subunit IFT machinery, HEK293 cells were co-transfected with human LRRC56 tagged with V5 and the reference IFT protein IFT88 fused to eYFP^13^. Immunoprecipitation using an anti-GFP antibody co-precipitated both IFT88-eYFP and LRRC56-V5, supporting an association of LRRC56 with the IFT machinery (Figure 1D)

The functional role of LRRC56 was further investigated in *T. brucei*, an organism that possesses a 9+2 axoneme and is amenable to genetic manipulation^23,40^. The *T. brucei* LRRC56 orthologue (Tb427.10.15260) comprises 751 amino acids and shares 42% sequence identity in the conserved leucine-rich repeat (LRR) region (Figure S2).

Since *T. brucei* maintains its mature flagellum during formation of the new one, it is possible to monitor both maintenance and assembly in the same cell ^41^. YFP-tagged LRRC56 (YFP::LRCC56) was expressed in cells upon endogenous tagging and the protein was detected in the distal portion of the new flagellum, whereas the old flagellum lacked the protein (Figure 2A). Following division, cells with one flagellum displayed two different profiles: either a strong signal towards the distal end of the flagellum (Figure 2B) or no signal (not shown). The first scenario reflects daughter cells that inherited a new flagellum, while the latter reflects those daughter cells that inherited an old flagellum. This observation is consistent with LRRC56 being recruited during flagellar assembly before its removal during flagellum maturation, after cell division. Co-localisation with the IFT protein IFT172, but not with the axoneme marker mAb25 suggests that LRRC56 associates with IFT and not dynein arms. Many, but not all, IFT trains contained LRRC56, suggesting it is a cargo rather than a core component. The LRRC56 signal is lost upon detergent extraction of the cytoskeleton, which strips the membrane and IFT particles but not the dynein arms (data not shown), further supporting association with IFT.

**Figure 2.**
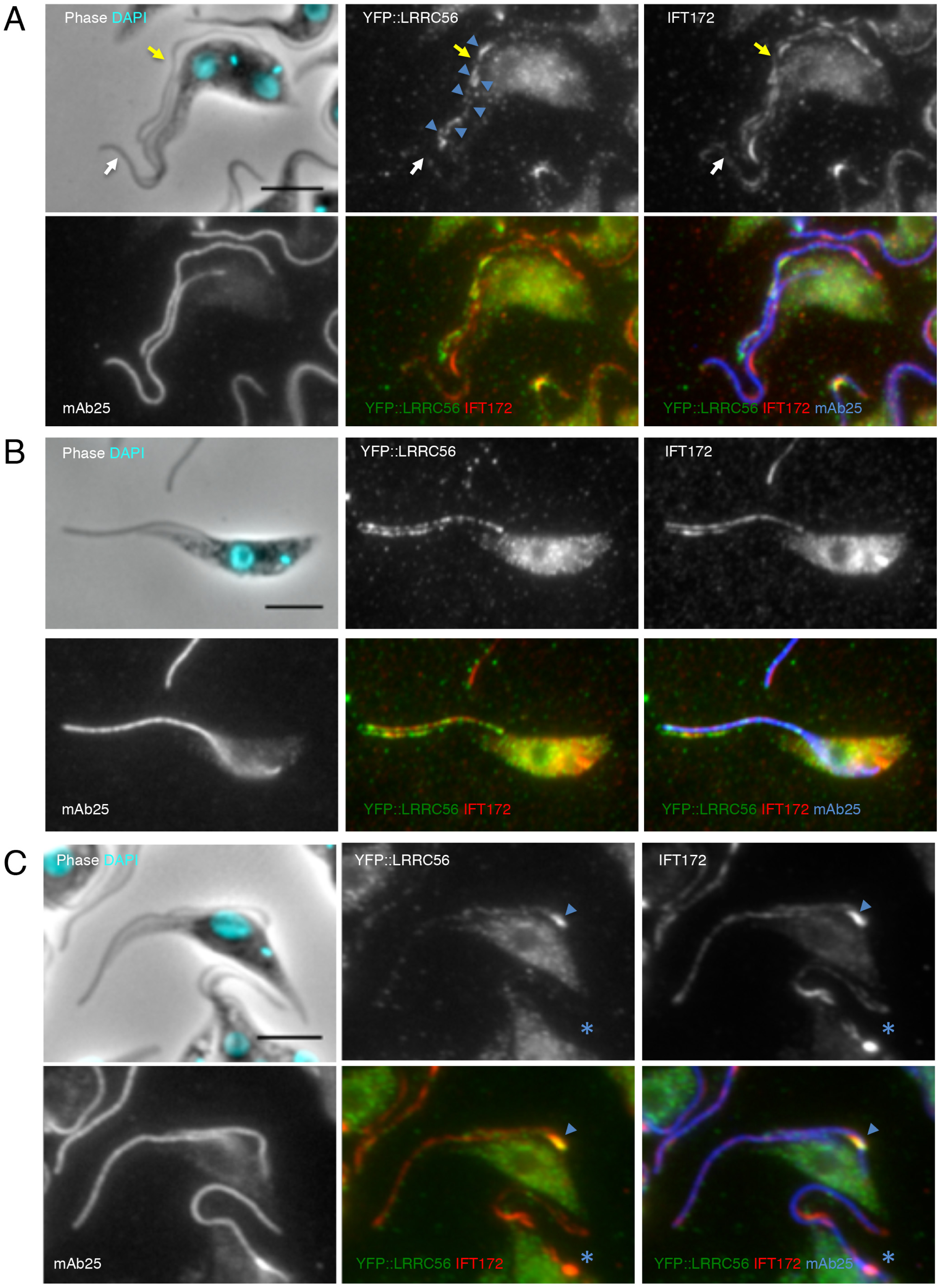
LRRC56 associates with IFT trains and not with the axoneme. An YFP::LRRC56 expressing cell that assembles its new flagellum (yellow arrow) shows staining (anti-GFP, green on merged images) in the new flagellum (blue arrowheads) that co-localises with the anti-IFT172 (red on merged images) but not with the anti-axoneme marker (mAb25, blue on merged images). No YFP staining is visible in the old flagellum (white arrow) whereas IFT172 positive trains are clearly present. IFT trains are predominantly found on microtubules doublets 4 and 7,^21^ hence the visual aspect of two separate tracks around the axoneme. DNA is stained with DAPI (cyan) showing the presence of two kinetoplasts (mitochondrial DNA) and two nuclei typical of cells at late stage of their cycle^41^. (B) Cytokinesis results in two daughter cells each containing a unique flagellum^41^. The one inheriting the new flagellum remains positive for YFP::LRRC56 that still shows association with IFT trains (same staining as in A). (C) The same immunofluorescence assay was performed on *IFT140^RNAi^* cells expressing YFP::LRRC56 after 24h in RNAi conditions. DNA staining shows that the top cell is mitotic and assembles a new flagellum. The IFT172 staining reveals the presence of a stalled IFT train that contains a considerable quantity of YFP::LRRC56 (arrowhead). The bottom cell is at a further stage of its cell cycle, yet its new flagellum is much shorter, indicating that RNAi knockdown occurred at a very early phase of construction (star). In these conditions, IFT172 staining shows that the very short flagellum contains a large amount of accumulated IFT material, yet no signal is visible for YFP::LRRC56 (star).

To further investigate the link with IFT, the distribution of YFP::LRRC56 was studied in trypanosome mutant cell lines in which tetracycline-inducible RNAi knockdown of an IFT-B (*IFT88^RNAi^*) or an IFT-A (*IFT140^RNAi^*) member results in either defective anterograde or retrograde transport, respectively^20,21^. In *IFT88^RNAi^* cells, the flagellar YFP::LRRC56 signal disappeared when anterograde transport was disrupted and the protein was found predominantly in the cytoplasm (Figure S3A and S3B). In the retrograde mutant *IFT140^RNAi^*, trains travel into the new flagellum but fail to be recycled to the base. The YFP::LRRC56 distribution pattern turned out to be more complex (Figure S3C and S3D) and required more detailed investigation. Cultures are not synchronised and RNAi knockdown can impact cells at different stages of flagellum construction. If this happened when flagella started to grow, IFT proteins accumulated in very short flagella, but LRRC56 was rarely detected (Figure 2C). However, when IFT arrest took place at later stages of elongation, LRRC56 was frequently found in high concentration always associated with stalled IFT trains (Figure 2C). These results reflect the association of LRRC56 as cargo of IFT trains, with a progressive increase during flagellar construction. After 2 days of tetracycline-inducible RNAi knockdown, the LRRC56 is no longer detected in the flagellum (Figure S3A and S3B).

LRRC56 localisation was unchanged in *DNAI1^RNAi^* and *ODA7 ^RNAi^* mutants that are defective in their dynein arm constitution or cytoplasmic preassembly, respectively (Figure S4). This shows that LRRC56 does not require the presence of dynein arms to be associated with flagella. Together, these findings reveal that LRRC56 likely performs an IFT-associated function in motile cilia.

### Absence or mutation of *T. brucei* LRRC56 is responsible for motility defects explained by absence of outer dynein arms in the distal segment of the axoneme

We next chose to assess any impact caused by the absence, or mutation of LRRC56. An *LRCC56* null mutant was generated by double gene knockout (Figure S5A), and whole-genome sequencing demonstrated that the *LRRC56* gene had been replaced by the drug resistance cassettes (Figure S5B). Visual examination showed a significant reduction in flagellar beating and cell swimming, with ∼10% of cells struggling to complete cell division and remaining attached by their posterior extremities (Figure S5C). These phenotypes indicate slow cytokinesis and increased generation times which, in trypanosomes, is typical of motility defects (Figure S5D)^22,42,43^. Microscopy revealed that *lrrc56-/-* cell motility is characterised by an erratic swimming pattern with altered propagation of the tip-to-base wave, increased frequency of base-to-tip waves, and frequent tumbling typical of outer dynein arm mutants (Figure 3) ^22,44^. Examination of the axoneme structure by TEM revealed that more than one third of sections analysed from *lrrc56-/-* cells showed absence of 6 to 9 ODAs (Figure 4). This was found in the distal portions of both old and new flagella, as indicated by the presence of a smaller cell body (or no cell body) ^45^. This suggests a role for LRRC56 in the assembly of dynein arms in the distal portion of the axoneme. Immunostaining with an anti-DNAI1 (dynein arm intermediate chain 1 also known as IC78) antibody^22,23^ that recognises an essential structural component of the ODA, confirmed that only the proximal axonemal region stained positively in *lrrc56* −/− cells (Figure 4H). Control cells stained from the base to the tip of all axonemes (Figure 4G), confirming that LRRC56 is essential for distal assembly of ODA.

**Figure 3.**
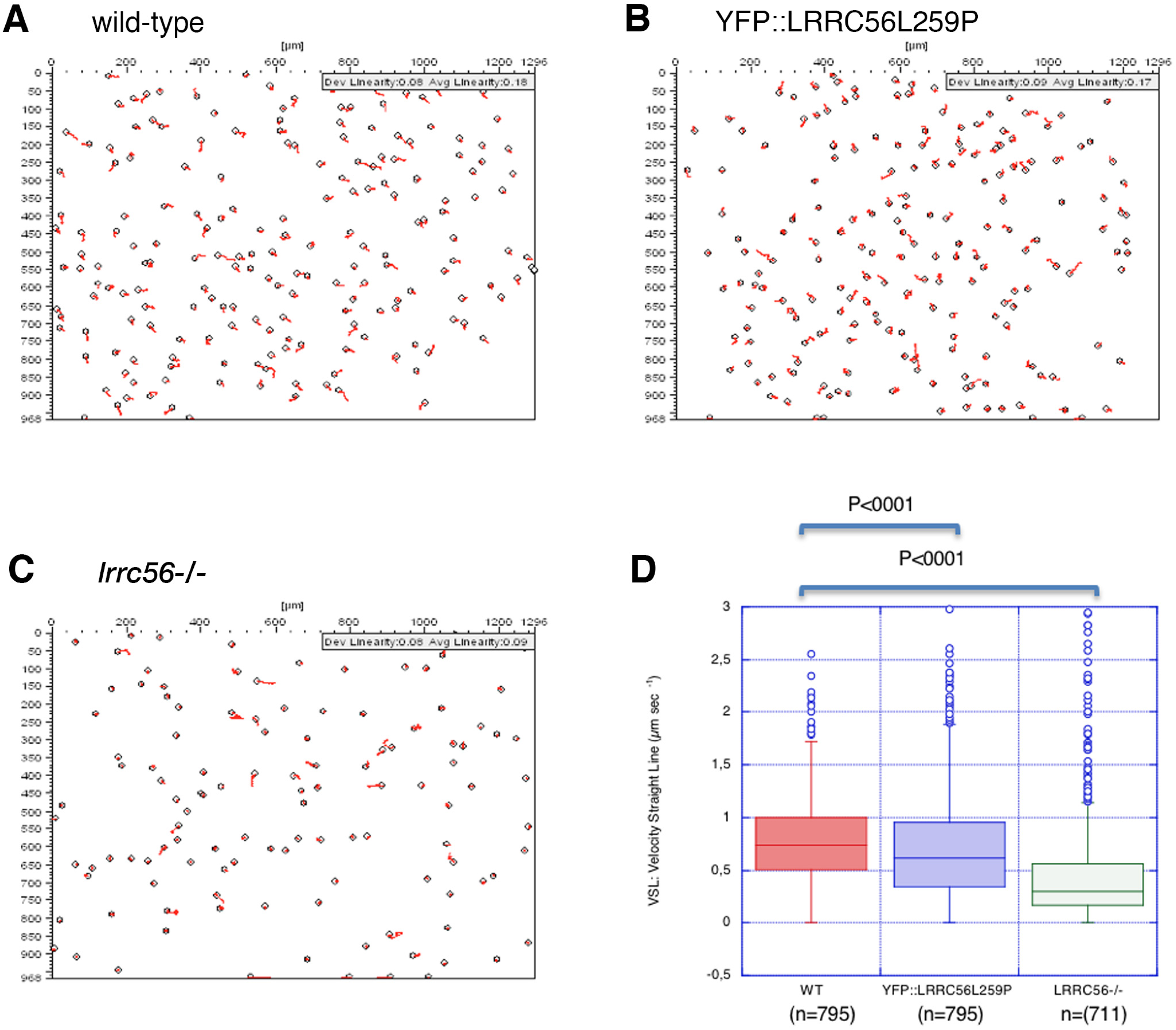
Absence of LRRC56 or expression of the L259P mutation reduces flagellum beating and cell motility. (A-C) Tracking analysis^34^ showing the movement of individual trypanosomes in wild-type control (A), in cells expressing only YFP::LRRC56L259P (B) and in *lrrc56-/-* cells (C). Sustained motility is only observed in control conditions. (D) Quantification of the straight-line movement confirms the visual impression that motility was reduced in a statistically significant manner in YFP::LRRC56L259P cells and almost abolished in *lrrc56-/-*. Total number of cells, mean and standard deviation are indicated on the figure. Statistical analysis was performed using t test.

**Figure 4.**
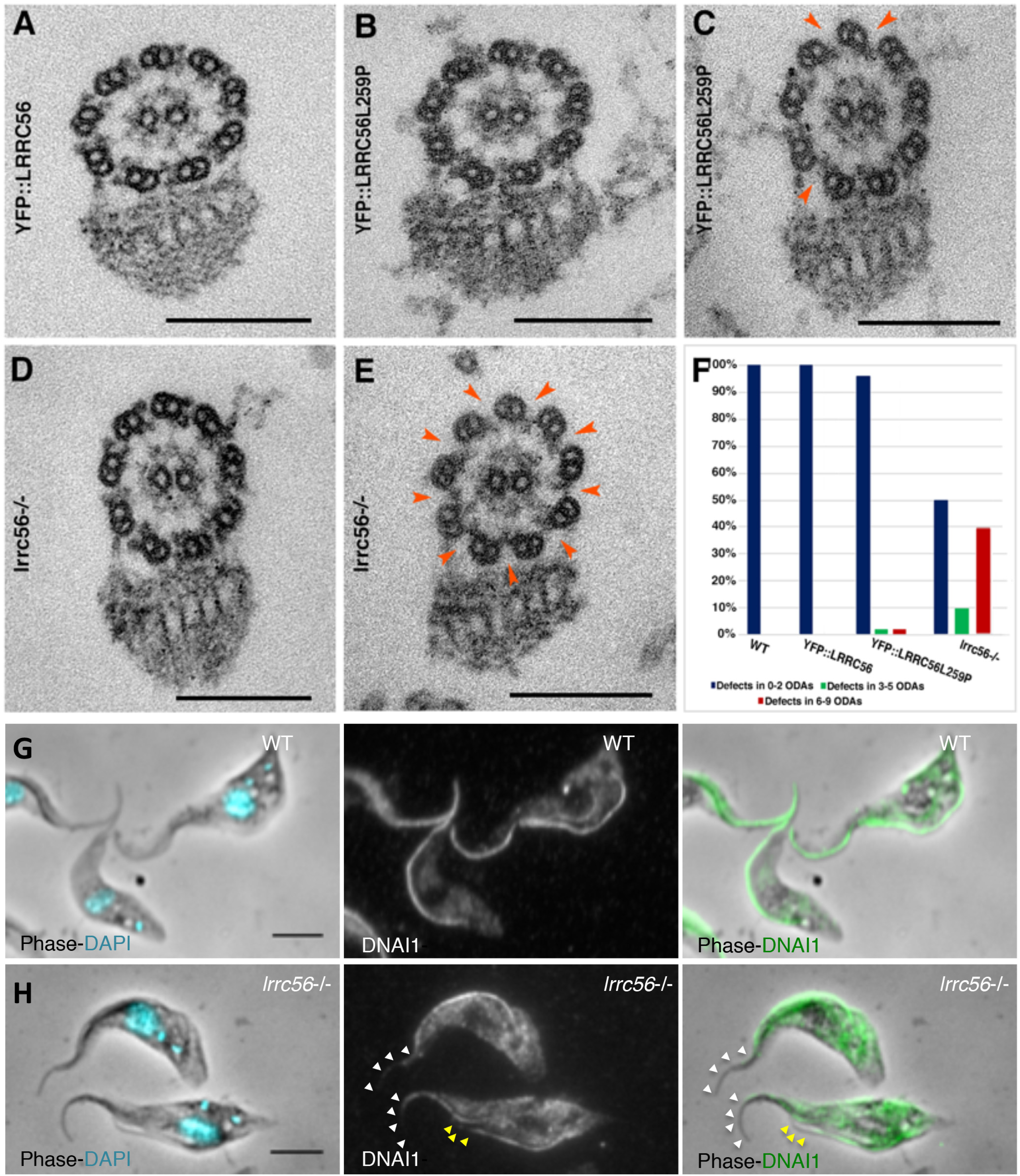
LRRC56 is required for assembly of ODA in the distal portion of the axoneme. (A-E) Sections are shown through the flagella of detergent-extracted cytoskeletons from various cell lines. Stripping the flagellum membrane and matrix facilitates the analysis of structures^22,52,53^. Sections through control YFP::LRRC56-expressing cells (A) possess all 9 ODA, whereas a mixture of sections with normal profiles or with several missing ODA (orange arrowheads) is encountered in YFP::LRRC56L259P-expressing cells (B-C) or *lrrc56-/-* cells (D-E). (F) Sections were grouped in three categories: defects in 2 ODA or less (blue), in 5 ODA or less (green) and in 6 ODA or more (red). The total counted number of sections is 50 for each sample. Full details are given in Table S2. (G-H) IFA with the anti-DNAI1 antibody stains the whole axoneme of wild-type cells (G) as expected^23^. However, the staining was limited to the proximal portion in *lrrc56-/-* cells in both growing (missing portion shown by yellow arrowheads) and mature flagella (white arrowheads) (H).

We next evaluated in trypanosomes the impact of the p.Leu259Pro mutation which corresponds to the p.Leu140Pro observed in Family 2. One *LRRC56* allele was deleted and the other allele was replaced endogenously with a L259P modification in YFP-tagged LRRC56 (YFP::LRRC56L259P cells). The YFP::LRRC56L259P protein localises normally to the distal portion of the new flagellum, showing that this residue is not required for proper expression and localisation of LRRC56 (data not shown). However, cells showed reduced motility, albeit not to the same extent as the double knockout (Figure 3C,D). Ultra-structural analysis of YFP::LRRC56L259P cells by TEM revealed a reduction in the number of ODAs, but that was less frequent compared to *lrrc56-/-* flagella (Figure 4B,C,F). We conclude that the mutated protein can associate with IFT trains, but functions less efficiently that its normal counterpart in supporting dynein arm assembly.

## DISCUSSION

Whole exome sequencing in two unrelated consanguineous families enabled us to identify homozygous variants in the *LRRC56* gene. The common clinical phenotype in the two families was cardiac laterality defects, while in Family 1 the affected individual also presented with recurrent pulmonary infections, and on investigation was found to have bronchiectasis. This combination of clinical features was suggestive of PCD, but a uniformly dyskinetic beat pattern was not observed. Investigation of cultured nasal epithelial cells from the affected individual in family 1 revealed a dyskinetic ciliary beat pattern, but no structural ciliary defects. We have observed this phenomenon before in our analysis of samples sent for the investigation of PCD (R.Hirst, personal communication).

In Family 1, with a homozygous splicing variant in LRRC56, further analysis confirmed that the mutated transcript is aberrantly spliced and is not predicted to encode a functional protein. In Family 2, a homozygous missense variant was identified. Functional investigation demonstrated a role for LRRC56 in the assembly of ODA in the protist *T. brucei.* We then modelled the effect of a null variant and the homozygous missense variant in this organism. We demonstrate that absence of LRRC56 or the presence of the homozygous missense variant seen in Family 2, both result in motility defects, caused by loss of ODAs in the distal segment of the axoneme. This observation is consistent with our interrogation of the role of LRRC56 in this organism.

Previous studies have shown that LRRC56 is found only in species with motile cilia that rely on IFT for axonemal assembly^39^. The reported biochemical distribution of LRRC56 is similar to that of IFT subunits, with about 50% of the protein associated with the membrane and matrix fraction^39^. Our pull-down experiments carried out in human cells expressing tagged LRRC56 and IFT88 revealed that LRRC56 interacts with the IFT machinery. In *T. brucei*, LRRC56 is recruited to the flagellar compartment at advanced stages of construction and co-localises with many (but not all) IFT trains. This flagellar localisation is dependent on IFT, as confirmed by analysis of cell lines defective in either anterograde or retrograde IFT. These results support a model in which LRRC56 associates with IFT trains and may function as a cargo adaptor to transport dynein arms towards the distal end of cilia and flagella.

The absence of LRRC56 has an impact on ciliary motility, as observed here in cultured cells from one affected patient and in *T. brucei*. This is consistent with the *oda8* (*LRRC56* gene homologue) null mutant in *Chlamydomonas*, although the impact on the presence of dynein arms was remarkably variable. ODAs are absent throughout the length of the flagellum in *Chlamydomonas* ^46^ whereas they are missing at only the distal part of the *T. brucei* flagellum. In both cases, this loss of ODA interferes with proper initiation of flagellum beating resulting in reduced motility and dyskinetic flagellar beating. In samples from the only human patient that were available for analysis, sections were analysed in different segments of the cilia (proximal, intermediate, distal), but dynein arms were not visibly affected.

Although the central core of the protein containing the leucine-rich repeat domains is well conserved, other segments are variable across species, with large and often unique N- and C-terminal extensions. This may translate into a common central function, such as association of LRRC56 with the IFT machinery for dynein arm transport, but could also result in considerable structural variation, potentially supporting species-specific interaction partners. Indeed, the composition of dynein arms (number and type of heavy, intermediate and light dynein chains) varies between species, as demonstrated by both biochemical and genetic analysis^42^. The outer dynein arm composition differs between the *Chlamydomonas* ODAs that contain 3 heavy dynein chains chains (α, β and γ), and the human and trypanosome ODA which each contain only 2 heavy chains (α and β).

In *Chlamydomonas*, early flagellum growth is very rapid (up to 10 µm per hour)^47^, and might necessitate a greater contribution from ODA8 to ODA transport compared to the slower growth rate observed in trypanosomes^48^, and respiratory epithelial cells ^49^. This could explain the absence of ODA throughout the axoneme compared to only the distal part in trypanosomes. We cannot exclude the possibility that discrete structural defects of dynein arms occur in the human patient whose cilia were analysed. For example, *DNAH11* nonsense mutations are associated with subtle dynein arm modifications that can only be detected by advanced TEM imaging and tomography ^3^. In this regard, approximately 30% of PCD patients showing ciliary dysfunction have no or very subtle ciliary ultrastructure abnormalities, when investigated with standard transmission electron microscopy^50^

Overall, our findings underline the evolutionary complexity of outer dynein arms assembly and trafficking. We have shown that bi-allelic mutations in the *LRRC56* gene are responsible for laterality defect in two unrelated families. We provide evidence of PCD-like symptoms caused by mutations in a gene encoding a protein required for ODA transport, rather than composition or assembly. Our work expands the list of ciliary genes involved in human disorders^51^, while also providing insight into the role of LRRC56 and cilia biology in human development.

## SUPPLEMENTAL DATA

Supplementary data consists of six figures, 4 videos and one table

## ACKNOWLEDGEMENTS

We wish to thank the families reported here for their willingness to participate in these research efforts. We thank the Ultrastructural BioImaging Plateforme for providing access to the TEM equipment. We thank Bruno Louis and Jean-François Papon, Institut National de la Santé et de la Recherche Médicale, U955, Créteil, France, for help in video-microscopy. Research at the Institut Pasteur is funded by an ANR grant (ANR-14-CE35-0009-01), by a French Government Investissement d’Avenir programme, Laboratoire d’Excellence “Integrative Biology of Emerging Infectious Diseases” (ANR-10-LABX-62-IBEID) and by La Fondation pour la Recherche Médicale (Equipe FRM DEQ20150734356). This work was supported, in part, by the Care4Rare Canada Consortium funded by Genome Canada, the Canadian Institutes of Health Research (CIHR), the Ontario Genomics Institute, Ontario Research Fund, Genome Quebec, and Children’s Hospital of Eastern Ontario Foundation and the Canadian Rare Diseases: Models and Mechanisms Network funded by CIHR and Genome Canada. The authors wish to acknowledge the contribution of the high-throughput sequencing platform of the McGill University and Génome Québec Innovation Centre, Montréal, Canada. Research at the University of Leeds was supported by grants MR/M009084/1 and MR/L01629X/1 awarded by the UK Medical Research Council.

## WEB RESOURCES

AgileMultiIdeogram, http://dna.leeds.ac.uk/agile/AgileMultiIdeogram/

Burrows-Wheeler Aligner, http://bio-bwa.sourceforge.net/

CASA Tracking V.5.5, https://safe.nrao.edu/wiki/bin/view/Software/CASA/WebHome

CLUSTAL Omega, https://www.ebi.ac.uk/Tools/msa/clustalo/

dbSNP, http://www.ncbi.nlm.nih.gov/projects/SNP/

Exome Aggregation Consortium (ExAC) Browser, http://exac.broadinstitute.org/

Genome Analysis Toolkit (GATK), https://www.broadinstitute.org/gatk/

Human Protein Atlas, https://www.proteinatlas.org

ImageJ, https://imagej.nih.gov/ij/

NHLBI Exome Variant server, http://evs.gs.washington.edu/EVS/

OMIM, http://www.omim.org/

Picard, http://broadinstitute.github.io/picard/

Protein BLAST, https://blast.ncbi.nlm.nih.gov/Blast.cgi?PAGE=Proteins

SAMtools, http://www.htslib.org/

TriTrypDB: The Kinetoplastid Genomics Resource, http://tritrypdb.org/tritrypdb/

SIFT, http://sift.jcvi.org

POLYPHEN2, http://genetics.bwh.harvard.edu/pph2/

CADD, http://cadd.gs.washington.edu

## Permissions

See email below from RH Hirst re personal communication

## Supplemental Data

**Figure S1.**
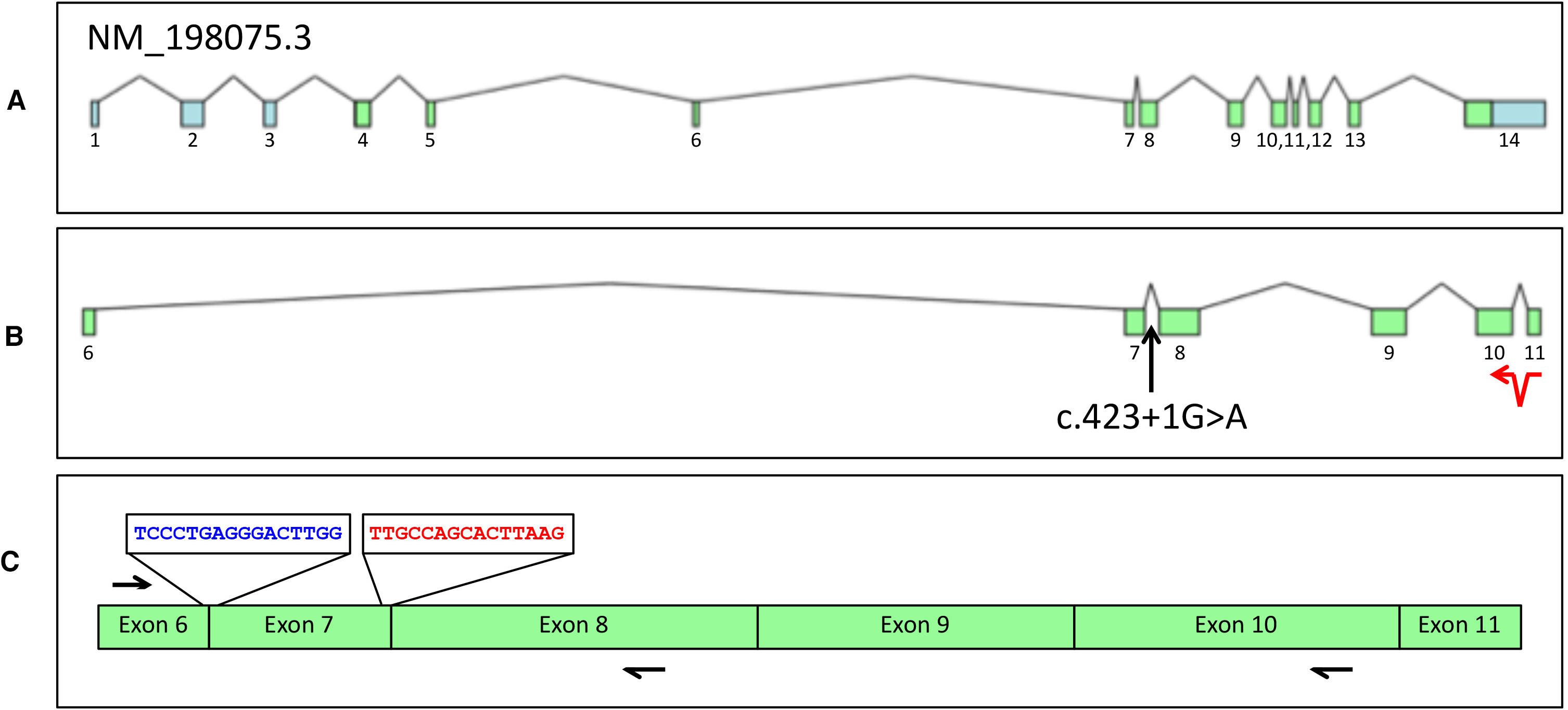
Primer locations for the *in vitro* splicing assay. (**A**). The *LRRC56* gene structure drawn to-scale for transcript NM_198075.3. (**B**) The location of the gene-specific primer (red arrow) used to perform the first-strand cDNA synthesis for the *in vitro* splicing assay for Family 1 individual IV:1. (C) Black arrows show the location of the exon 6 forward primer used in conjunction with either exon 8 or exon 10 reverse primers. Blue text represents the string used to extract *LRRC56*-specific reads spanning the exon 6/7 junction. The 2 nucleotides located beyond the red text in each extracted sequence read were interrogated to determine whether the read represent a spliced or unspliced transcript. At least 6.9 million reads were generated per primer combination per sample. Only a small proportion of these were LRRC56-specific, with the 6F/8R primer combination working more effectively than 6F/10R. For the control sample, these represented 0.46% and 0.08% of total reads respectively. The nucleotides located beyond the 3′ end of exon 7 were assayed for the correctly spliced exon 8 sequence (GA) and the ratio of spliced to unspliced transcripts was calculated. In the control sample, 98.4% and 99.9% of detected transcripts were correctly spliced, suggesting the assay was biologically representative. In contrast, 100.0% and 89.16% of reads from the affected subject were unspliced.

**Figure S2.**
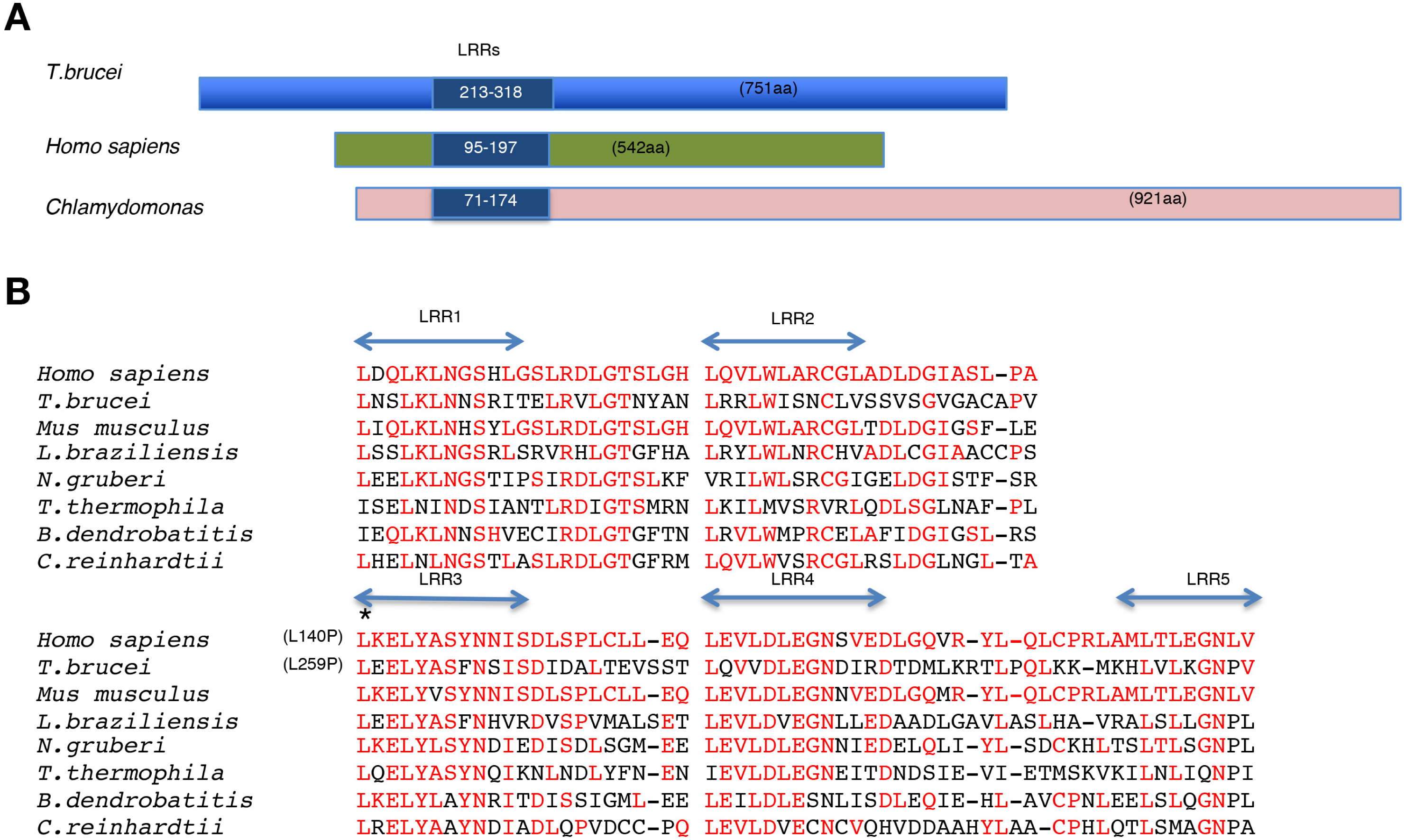
LRRC56 is conserved in ciliated organisms. (**A**) Alignment of LRCC56 predicted protein sequences from *T. brucei*, *H. sapiens* and *C. reinhardtii*. While the leucine-rich domain is evolutionary conserved, the flanking portions are divergent and vary in length between species. (**B**) Alignment of the leucine-rich domain in *H. sapiens, T. brucei* and three neighbouring species (*Trypanosoma congolense*, *Trypanosoma cruzi* and *Leishmania braziliensis*), *C. reinhardtii* and *M. musculus*. Blue arrows indicate the 5 leucine-rich repeat motifs. Conserved residues are highlighted in red. A star (*) marks the position of the human L140P mutation; the equivalent residue in *T. brucei* is located at position 259 due to the N-terminal extension in this organism.

**Figure S3.**
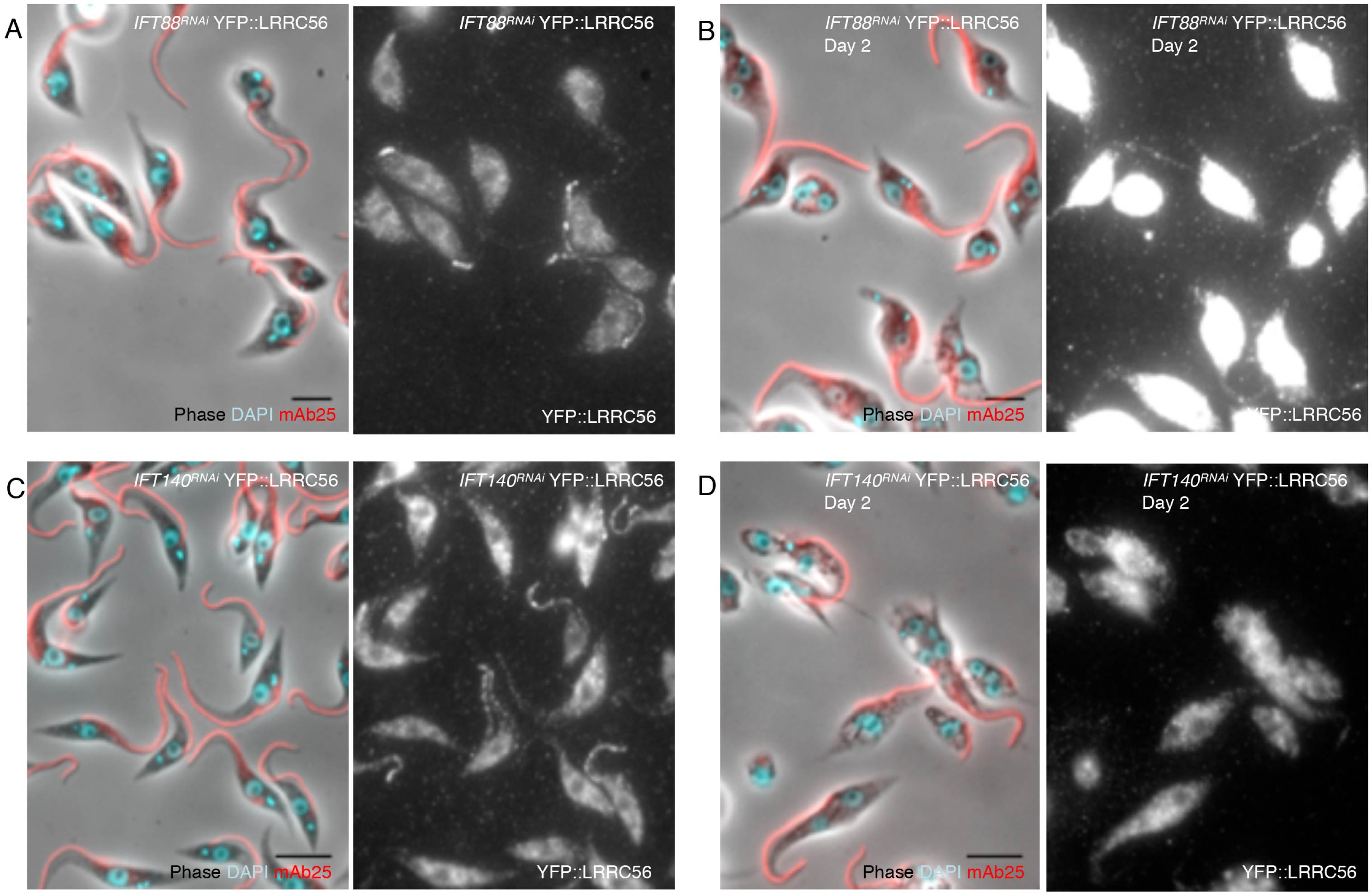
Inhibition of IFT anterograde transport prevents flagellum targeting of YFP::LRRC56. Upon endogenous tagging, YFP::LRRC56 was expressed in *IFT88^RNAi^* cells or *IFT140^RNAi^* cells without induction (**A,C**) or with (**B,D**) 48h-induction of RNAi. Whereas in non-induced conditions the usual pattern of YFP::LRRC56 staining associated with IFT trains in both cell types is observed in the growing flagellum or in the daughter flagellum upon cell division (A,C), the staining is virtually absent in flagella of *IFT88^RNAi^* induced cells and the protein is redistributed to the cytoplasm (B). In induced *IFT140^RNAi^* cells, the LRRC56 is no longer detected in the flagellum (D). For each sample, left panels show phase contrast images with DAPI (blue) and axonemal staining (mAb25, red) whereas right panels show the immunofluorescence assay staining of YFP::LRRC56 with anti-GFP.

**Figure S4.**
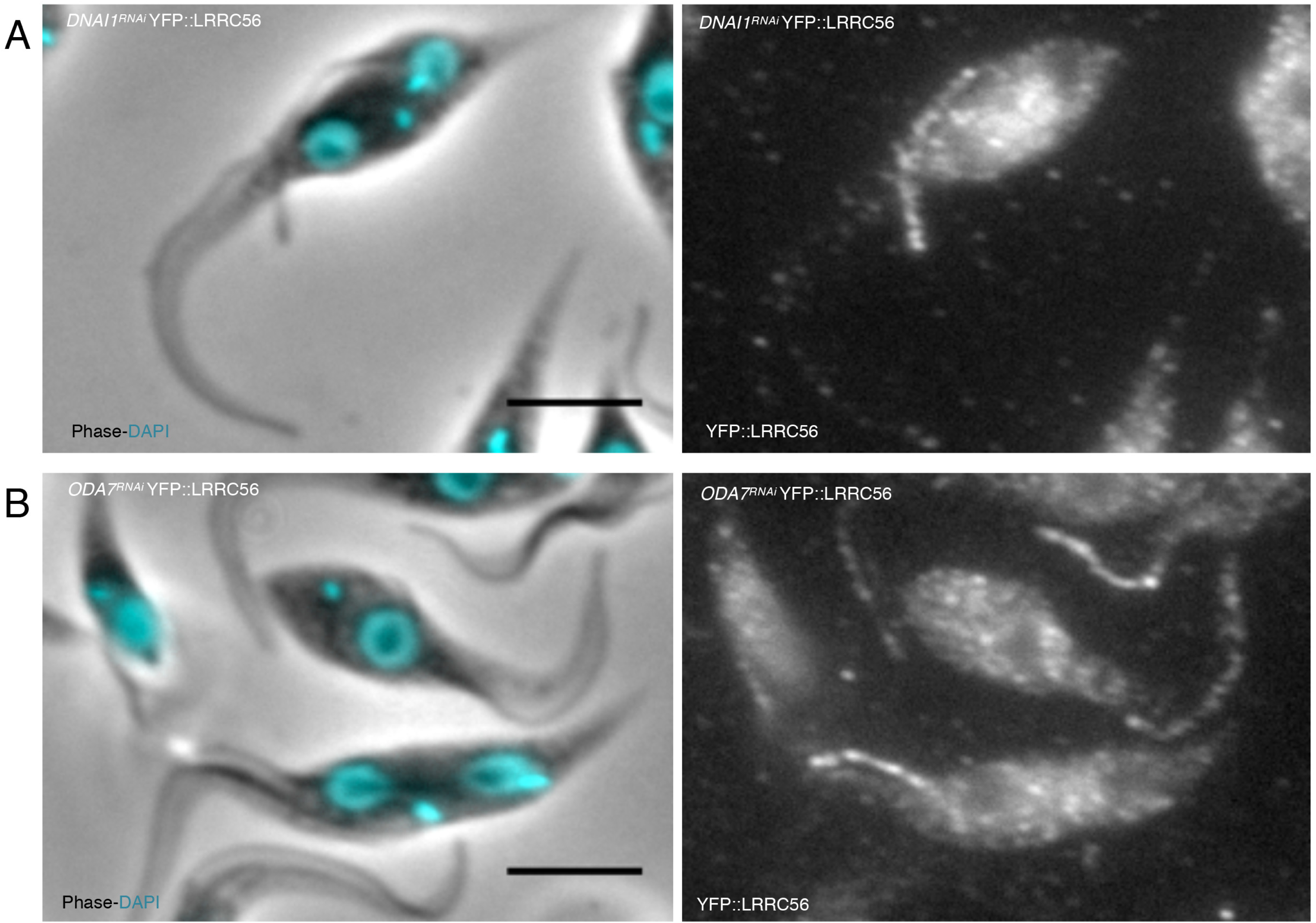
LRRC56 association with IFT trains is independent of the presence of ODA. Upon endogenous tagging, YFP::LRRC56 was expressed in *DNAI1^RNAi^* (**A**) or *ODA7^RNAi^* cells. DNAI1 (or IC78) is a dynein intermediate chain and an essential component of the ODA^22^, and ODA7 (or LRRC50) is required for cytoplasmic pre-assembly of the ODA^23^. The YFP::LRRC56 distribution is identical in non-induced (not shown) and induced conditions for both *DNAI1^RNAi^* (**A**) or *ODA7^RNAi^* (**B**) cells, showing that flagellum targeting of LRRC56 and its association to IFT trains is independent of the presence of complete ODA. Left panels show phase contrast images with DAPI (blue) and right panels show the staining of YFP::LRRC56 with the anti-GFP.

**Figure S5.**
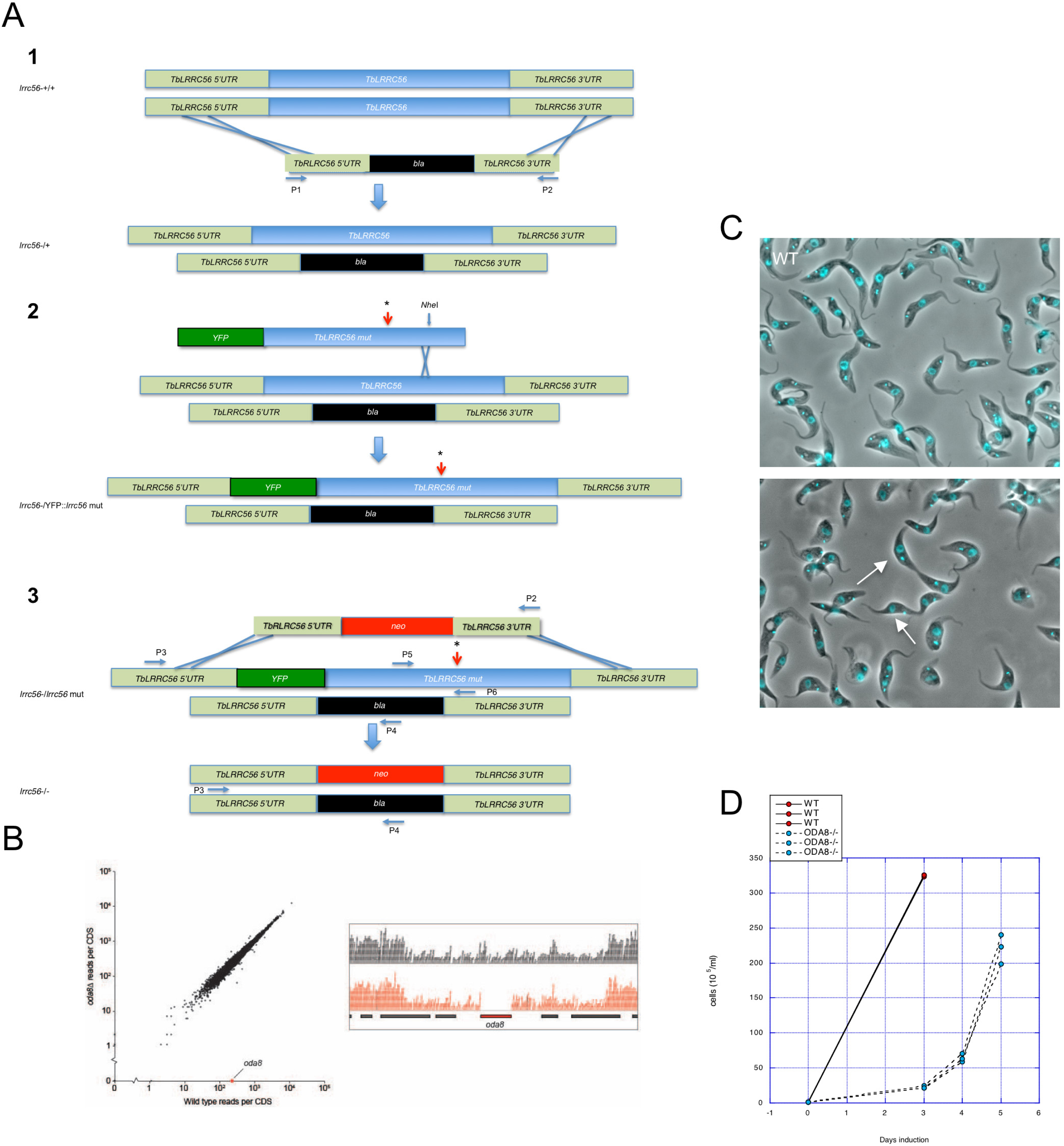
Double gene knockout of *LRRC56* in *T. brucei* affects cell motility, cell division and cell growth. (**A**) Summary of the strategy used for endogenous tagging and for double gene deletion of *LRRC56* in *T. brucei*. 1. One *LRRC56* allele was replaced with a blasticidin resistance marker. 2. The remaining *LRRC56* allele was replaced with the mutated *YFP::LRRC56L259P* gene in this single knockout cell line. 3. This tagged allele was replaced with a G418 drug resistance marker. (**B**) Next generation sequencing confirmed *lrrc56* knockout in *lrrc56-/-* strain. Left, scatter plot of non-redundant gene set comparing reads per coding sequence in wild type and *lrrc56-/-*strains. Right, visualisation of reads at the *LRRC56* locus on chromosome 10. Black reads (top track): wild type, red reads: *lrrc56-/-*. (**C**) Phase contrast image of fields of cells stained with DAPI (cyan) in wild-type (WT) or *lrrc56-/-* cultures. The increased presence of cells at late phase of cytokinesis (white arrows) is visible. This is explained by the role of motility to complete cell division and to allow separation of daughter cells ^4,22,43^. (**D**) Growth was monitored over 5 days in wild-type cells and *lrrc56-/-* cells. Triplicates are shown.

**Figure S6.**
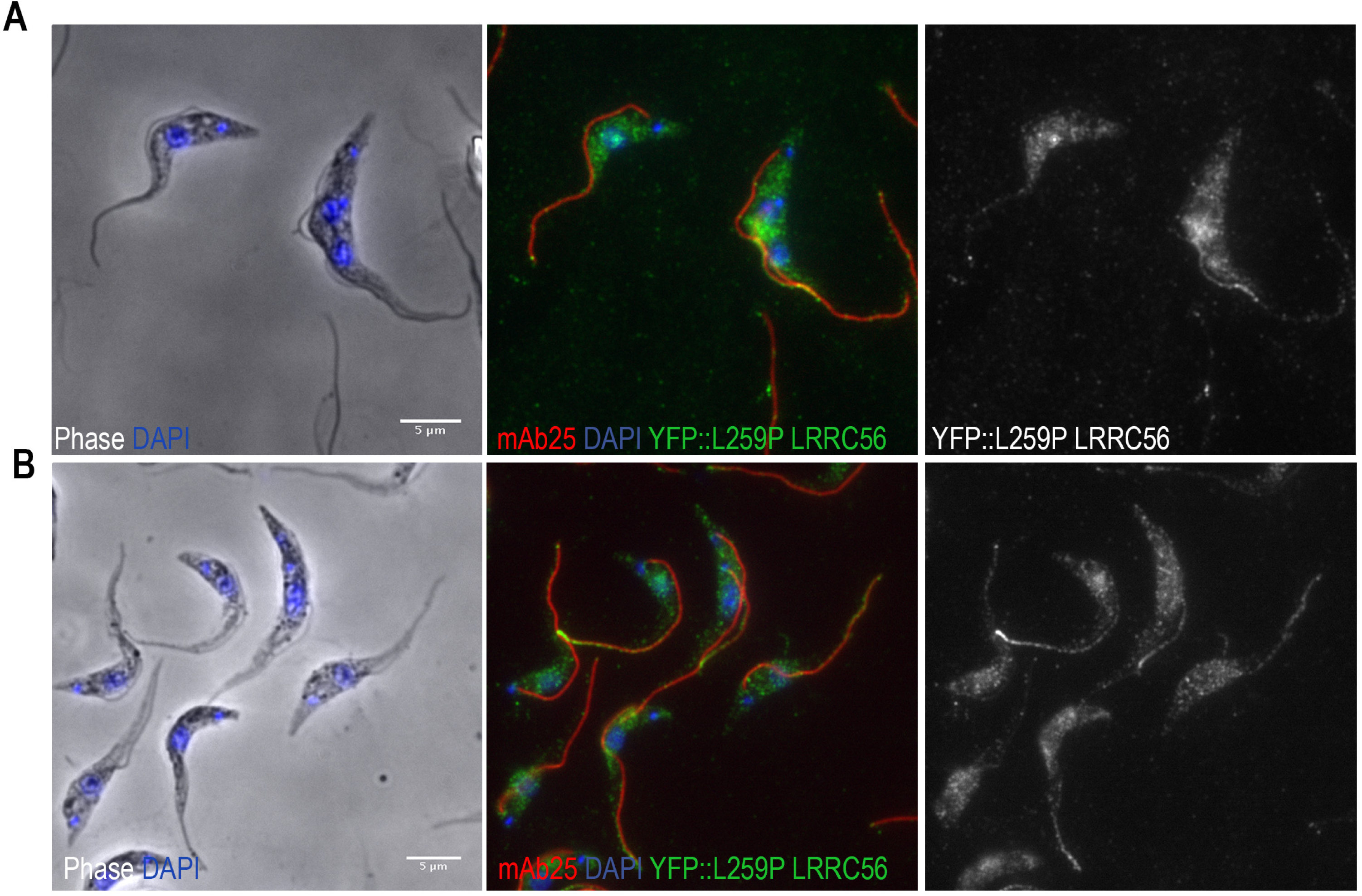
Expression of YFP::LRRC56 bearing the L259P mutation. Localisation by IFA of LRRC56-/YFP::LRRC56L259P in *T. brucei* cells obtained using strategy depicted Figure S5A (steps 1-2). (A & D) methanol-fixed cells stained with anti GFP or anti-axoneme mAb25 antibody. Left panels show phase contrast images with DAPI (blue), mid panels are images overlays obtained for GFP (green) mAb25 (red) and DAPI) (blue) signals and right panels show the IFA staining of YFP::LRRC56L259P with the anti-GFP (white).

**Table S1.** Genomic coordinates of the MultiIdeogram identified autozygous intervals. The shared region on chromosome 11 which contains *LRRC56* is highlighted in bold

| Individual | Chromosome | Start | End | Length (bp) |
| --- | --- | --- | --- | --- |
| Family 1, IV:1 | 1 | 16,532,803 | 17,023,209 | 490,406 |
| Family 1, IV:1 | 1 | 24,194,748 | 38,006,359 | 13,811,611 |
| Family 1, IV:1 | 1 | 94,511,533 | 101,203,827 | 6,692,294 |
| Family 1, IV:1 | 1 | 117,487,710 | 120,611,960 | 3,124,250 |
| Family 1, IV:1 | 1 | 145,112,414 | 147,381,397 | 2,268,983 |
| Family 1, IV:1 | 1 | 149,905,279 | 211,280,724 | 61,375,445 |
| Family 1, IV:1 | 1 | 222,798,106 | 241,712,134 | 18,914,028 |
| Family 1, IV:1 | 1 | 248,084,511 | 248,525,071 | 440,560 |
| Family 1, IV:1 | 2 | 130,832,292 | 131,674,285 | 841,993 |
| Family 1, IV:1 | 2 | 197,118,729 | 207,621,759 | 10,503,030 |
| Family 1, IV:1 | 2 | 220,072,766 | 220,421,417 | 348,651 |
| Family 1, IV:1 | 2 | 228,111,435 | 231,407,663 | 3,296,228 |
| Family 1, IV:1 | 3 | 49,739,507 | 52,256,697 | 2,517,190 |
| Family 1, IV:1 | 3 | 105,260,596 | 113,858,350 | 8,597,754 |
| Family 1, IV:1 | 3 | 195,511,556 | 195,518,112 | 6,556 |
| Family 1, IV:1 | 4 | 106,474,096 | 114,177,035 | 7,702,939 |
| Family 1, IV:1 | 4 | 138,449,683 | 155,407,680 | 16,957,997 |
| Family 1, IV:1 | 5 | 73,930,751 | 75,427,935 | 1,497,184 |
| Family 1, IV:1 | 5 | 122,685,801 | 137,507,104 | 14,821,303 |
| Family 1, IV:1 | 5 | 140,228,164 | 140,531,746 | 303,582 |
| Family 1, IV:1 | 6 | 29,855,570 | 33,382,288 | 3,526,718 |
| Family 1, IV:1 | 6 | 49,701,523 | 96,651,740 | 46,950,217 |
| Family 1, IV:1 | 7 | 27,135,314 | 36,320,821 | 9,185,507 |
| Family 1, IV:1 | 7 | 83,631,376 | 138,455,988 | 54,824,612 |
| Family 1, IV:1 | 9 | 125,486,968 | 127,228,596 | 1,741,628 |
| Family 1, IV:1 | 9 | 130,457,332 | 131,898,880 | 1,441,548 |
| Family 1, IV:1 | 10 | 112,360,315 | 118,383,463 | 6,023,148 |
| Family 1, IV:1 | 10 | 120,919,246 | 135,491,083 | 14,571,837 |
| **Family 1, IV:1** | **11** | **193,863** | **1,643,228** | **1,449,365** |
| Family 1, IV:1 | 11 | 56,143,394 | 56,143,785 | 391 |
| Family 1, IV:1 | 11 | 62,292,882 | 78,516,318 | 16,223,436 |
| Family 1, IV:1 | 11 | 95,825,374 | 110,023,628 | 14,198,254 |
| Family 1, IV:1 | 12 | 10,137,557 | 11,183,451 | 1,045,894 |
| Family 1, IV:1 | 12 | 52,284,233 | 57,619,209 | 5,334,976 |
| Family 1, IV:1 | 12 | 123,200,247 | 124,325,977 | 1,125,730 |
| Family 1, IV:1 | 13 | 92,345,579 | 102,106,276 | 9,760,697 |
| Family 1, IV:1 | 14 | 24,429,093 | 30,066,976 | 5,637,883 |
| Family 1, IV:1 | 14 | 104,573,690 | 105,411,781 | 838,091 |
| Family 1, IV:1 | 15 | 52,404,862 | 91,524,841 | 39,119,979 |
| Family 1, IV:1 | 16 | 88,134,285 | 88,872,145 | 737,860 |
| Family 1, IV:1 | 17 | 6,115 | 970,413 | 964,298 |
| Family 1, IV:1 | 17 | 39,081,713 | 39,165,174 | 83,461 |
| Family 1, IV:1 | 18 | 44,635,048 | 60,036,083 | 15,401,035 |
| Family 1, IV:1 | 19 | 47,823,038 | 48,284,025 | 460,987 |
| Family 1, IV:1 | 19 | 57,175,484 | 57,909,872 | 734,388 |
| Family 1, IV:1 | 20 | 1,298,794 | 1,616,206 | 317,412 |
| Family 2, IV:3 | 1 | 22,141,206 | 22,846,709 | 705,503 |
| Family 2, IV:3 | 1 | 87,549,858 | 89,652,102 | 2,102,244 |
| Family 2, IV:3 | 1 | 147,121,977 | 151,396,037 | 4,274,060 |
| Family 2, IV:3 | 1 | 155,026,942 | 156,280,971 | 1,254,029 |
| Family 2, IV:3 | 1 | 162,344,390 | 169,272,453 | 6,928,063 |
| Family 2, IV:3 | 1 | 223,900,408 | 226,074,563 | 2,174,155 |
| Family 2, IV:3 | 1 | 247,921,100 | 248,487,768 | 566,668 |
| Family 2, IV:3 | 2 | 200,173,684 | 212,251,864 | 12,078,180 |
| Family 2, IV:3 | 2 | 220,239,561 | 220,435,034 | 195,473 |
| Family 2, IV:3 | 4 | 17,517,099 | 36,292,018 | 18,774,919 |
| Family 2, IV:3 | 4 | 10,716,8431 | 111,398,208 | 4,229,777 |
| Family 2, IV:3 | 5 | 15,858,8633 | 169,461,547 | 10,872,914 |
| Family 2, IV:3 | 6 | 28,228,342 | 33,382,288 | 5,153,946 |
| Family 2, IV:3 | 6 | 35,027,927 | 57,055,354 | 22,027,427 |
| Family 2, IV:3 | 6 | 62,284,247 | 79,595,167 | 17,310,920 |
| Family 2, IV:3 | 7 | 55,266,417 | 64,453,136 | 9,186,719 |
| Family 2, IV:3 | 7 | 147,926,734 | 151,818,777 | 3,892,043 |
| Family 2, IV:3 | 8 | 29,927,300 | 38,005,835 | 8,078,535 |
| Family 2, IV:3 | 8 | 63,978,658 | 92,136,728 | 28,158,070 |
| Family 2, IV:3 | 9 | 113,192,655 | 116,931,737 | 3,739,082 |
| Family 2, IV:3 | 9 | 127,101,924 | 129,102,840 | 2,000,916 |
| Family 2, IV:3 | 9 | 137,672,012 | 141,016,262 | 3,344,250 |
| **Family 2, IV:3** | **11** | **197,557** | **7,324,475** | **7,126,918** |
| Family 2, IV:3 | 11 | 56,143,198 | 56,143,823 | 625 |
| Family 2, IV:3 | 11 | 64,066,859 | 64,572,557 | 505,698 |
| Family 2, IV:3 | 12 | 10,165,469 | 12,618,715 | 2,453,246 |
| Family 2, IV:3 | 13 | 103,385,015 | 109,459,020 | 6,074,005 |
| Family 2, IV:3 | 16 | 16,138,313 | 33,961,370 | 17,823,057 |
| Family 2, IV:3 | 16 | 46,655,210 | 72,991,715 | 26,336,505 |
| Family 2, IV:3 | 16 | 89,167,075 | 89,597,221 | 430,146 |
| Family 2, IV:3 | 19 | 17,306,031 | 17,547,292 | 241,261 |
| Family 2, IV:3 | 20 | 32,295,541 | 33,764,632 | 1,469,091 |
| Family 2, IV:3 | 20 | 56,846,580 | 62,737,318 | 5,890,738 |

Positions reported for human genome build hg19.

